# Frontal midline theta power accounts for inter-individual differences in motor learning ability

**DOI:** 10.1101/2025.01.23.634624

**Authors:** Yuya Fukuda, Kazumasa Uehara

**Author notes:** Correspondence to: Kazumasa Uehara, Associate Professor, Ph.D. Department of Computer Science and Engineering, Toyohashi University of Technology Address: 1-1 Hibarigaoka, Tempaku-cho, Toyohashi, Aichi, JAPAN, Zip code: 441-8580.

## Abstract

Recent neurophysiological studies have demonstrated that frontal midline theta (FMT) activity plays a significant role in motor learning. One of the key challenges in motor learning is to understand the interindividual variability in learning proficiency rates, yet the underlying neural mechanisms remain unclear. To address this open question, this study recorded electroencephalogram activity during a visuomotor tracking task to investigate whether modulation of FMT power and the consistency of theta phase during motor preparation could explain individual differences in learning proficiency. We found a significant positive correlation between increased FMT power during motor preparation and learning proficiency rates. Specifically, individuals with greater FMT power exhibited faster learning rates. In contrast to this, no significant correlation was observed between the consistency of theta phase during motor preparation and learning proficiency. Together, these findings highlight that the FMT power, rather than phase synchrony, is closely associated with motor learning efficiency. This study provides a novel perspective for understanding the causes of individual differences in motor learning and further corroborates the previous evidence showing FMT power contributes to motor learning processes.

## Introduction

Humans possess the ability to adapt to diverse environmental changes and enhance their motor skills through learning. This ability is observed across a wide range of situations from the development to advanced musical performance or acquisition of sports skills. Moreover, many of the skill acquisition are characterized by their long-term retention (Park et al. 2013; Park and Sternad 2015). However, based on empirical evidence, the duration or repetition of practice required for the skill acquisition differ among individuals (Golenia et al. 2014). The open question here is how inter-individual variability of motor learning is defined neurophysiologically. Understanding these inter-individual differences is anticipated to provide new insights into the underlying processes of motor learning and performance (Seidler and Carson 2017; Anderson et al. 2021; Moore and Cluff 2021).

To tackle this, electroencephalogram (EEG), which is known to be a non-invasive method with high temporal resolution, would be helpful to investigate neural mechanisms underlying sensorimotor control. This is because EEG has clarified neural mechanisms underlying motor planning, preparation, and action, and post-movement processing (Reuter et al. 2022; Uehara et al. 2023b). Especially in the predefined EEG frequency bands such as delta, theta, alpha, beta, and gamma allows us to demonstrate neural correlations with cognitive and motor functions (Herrmann et al. 2005; Harmony 2013; Ramos-Murguialday and Birbaumer 2015; Struber et al. 2021; Beste et al. 2023).

Here we leveraged frontal midline theta (FMT, 4-8Hz) as a cue to investigate inter-individual differences in motor learning. Previous evidence suggests that FMT reflects cognitive control (Cavanagh and Frank 2014) and that this neural signal originates from the anterior cingulate cortex (ACC) (Mitchell et al. 2008; Womelsdorf et al. 2010). The ACC allocates cognitive resources to tasks requiring effort or conflict resolution, which is essential for adapting behavior in response to changes in the environment or task demands. The function of the ACC, which serves as proxy for FMT, also plays a central role in motor learning. An increase in FMT is significantly correlated with error correction through motor learning (Arrighi et al. 2016; Jonker et al. 2021). The degree of preparatory theta phase has been shown to highly predict perceptual performance (Busch et al. 2009; Busch and VanRullen 2010; Hanslmayr et al. 2013; Tomassini et al. 2017). Although we presume that FMT power and phase also contribute to motor learning proficiency, the extent to which they impact inter-individual variability of motor learning proficiency remains debatable.

Here we aimed to investigate the neural correlation between the modulation of FMT power during motor preparation and motor learning ability. A previous human EEG study reported that FMT power during movement planning increases with error reduction (Gentili et al. 2011). Based on this empirical finding, specially, we hypothesized that individuals who exhibit faster error reduction would show greater modulation of FMT power during motor preparation. Furthermore, this study discovered to comprehensively elucidate the interplay between FMT power and theta phase during motor preparation. This is because EEG power focuses more on the magnitude of local neural activity, whereas EEG phase indicates the timing within an oscillatory cycle, i.e., the temporal modulation of neural information processing. Addressing both neural features provides a clue to individual differences in motor learning ability.

## Materials and Methods

*Participants.* Twenty-five healthy volunteers (21 male, mean age = 21.76 ± 0.92 years, age range: 20-24 years) were participated in this study. All participants were native concerning our visuomotor learning task and had a normal or corrected-to-normal vision, no history of musculoskeletal or neurological disorders. Twenty-four out of 25 participants were right-handed and one participant was left-handed according to the Edinburgh Handedness Inventory (Oldfield 1971). All participants had a sufficient understanding of the experimental procedures and gave written informed consent prior to the data collection. This experimental protocol was approved by the Institutional Review Boards Involving Human Subjects of Toyohashi University of Technology in accordance with the guidelines established in the Declaration of Helsinki. Four participants were excluded from the final reports due to the following reasons: one participant was excluded due to technical issues with the data collection system, another participant due to failure to fit behavioral data during the analysis process, and two participants due to excessive EEG noises throughout the trials. We therefore reported the results obtained from twenty-one participants.

*Apparatus.* The present study employed a visuomotor learning task in which a cursor (yellow-closed circle) displayed on a personal computer (PC) screen was controlled as intended by generating forces with the right index and little fingertips and tracked accurately the target with the cursor (Fig.1A). The participants were required to press a force-torque transducer (USL06-H5, Tec Gihan, Co. Ltd., Japan) along the Z-axis to track a moving target (red-opened circle) in a straight line. These transducers were assigned to the index and little fingertips, respectively. Time series of force data were amplified through an amplifier device (GDA-06B, Tec Gihan, Co. Ltd., Japan) and then stored on a computer using a NI-DAQ (USB-6002, National Instruments, United States) via NI-DAQmx software at a sampling frequency of 1 kHz with analog-to-digital conversion. The index fingertip controls leftward cursor movement, while the little fingertip is responsible for rightward cursor movement. Therefore, the cursor direction was determined by the vector component calculated from the force applied by the index and little fingertips. Cursor velocity was also controllable by pressing hard or weakly the force-torque transducer. Visual cues, cursor control, cursor and target positions on the screen, and trigger signals sent to an EEG amplifier were managed by a dedicated software package (Graphical Design Lab, Japan), implemented in National Instruments LabVIEW software (version 2019 National Instruments, United States). We used a 24.5-inch PC monitor (XL2546K-B, BenQ; refresh rate of 144Hz), placed in front of a subject, to display visual cues, cursor, target, and binary visual feedback on each trial (*See the Experimental protocol and task section*). This monitor was positioned at a distance of 50 cm from the chin rest, aligned with the participant’s line of sight.

**Fig. 1.**
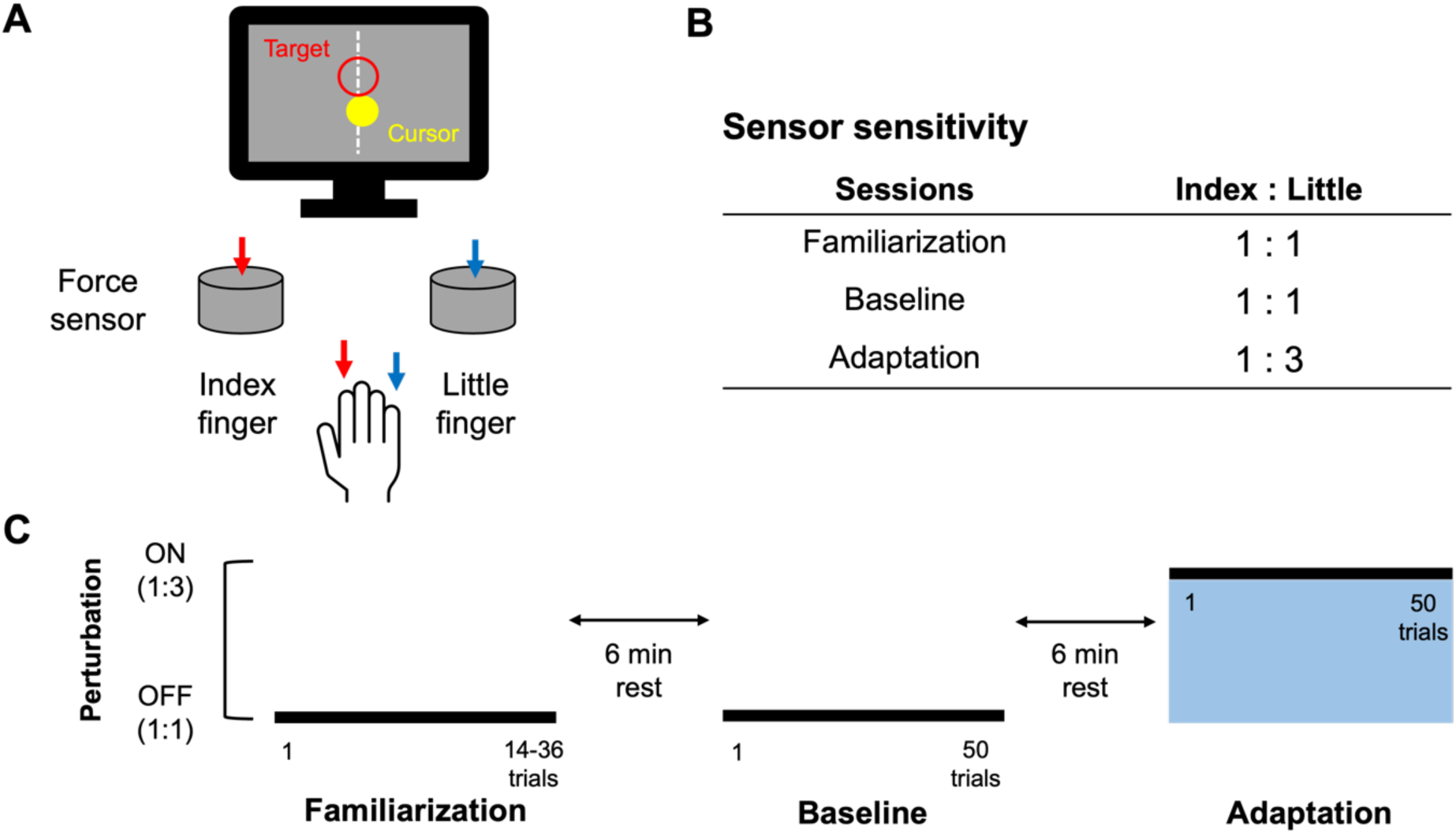
Experimental design and protocol. **(A)** Participants pushed the two force sensors using their index and little fingertips, respectively for controlling a cursor displayed on a PC monitor. **(B)** This experiment consisted of three sessions, and this table shows the sensitivity of the sensors in each session. **(C)** Data collection consisted of three blocks. The motor learning condition was made by changing the sensitivity of sensors for the little fingertips. The sensitivity of the sensor on the little fingertip was tripled from the learning session onward.

*Experimental protocol and task*. Apparatus used in this study is shown in Fig.1A. The experimental room was darkened to minimize the availability of any visual information that not projected on a PC monitor. Participants sat upright on a comfortable chair with their right forearm supported by a rigid table and their head stabilized using a chin rest to avoid involuntary muscle contractions. All participants underwent spontaneous EEG recordings in the resting state under both eyes-open and eyes-closed conditions before the visuomotor learning task. The eyes-closed condition required maintaining a resting state for 3 min without any thoughts or body movement. In the eyes open condition, subjects were instructed to view a video footage of natural scenery for 3 min without any thoughts or body movement. Naturalistic viewing is a potential alternative to traditional resting-state EEG recording, which typically involve watching a fixation cross. This protocol reduces eye movement-related artifacts and captures more naturalistic brain activity without causing arousal changes (Welke and Vessel 2022).

In this study, we asked participants to learn cursor control using force production with their right index and little fingertips. Fig.1C illustrates the experimental protocol and task structure. All subjects completed baseline and adaptation periods after task familiarization. Before starting the experimental task, we measured subject’s maximum muscle force production of the index and little fingertips of the right hand three times by pressing each sensor with maximal force. The sensor inputs were normalized by the obtained maximum muscle force. The cursor movement was defined according to the following formular:

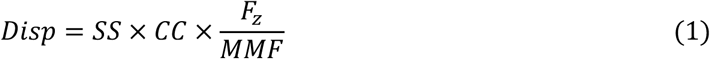

where *SS* the sensor sensitivity, *CC* is the correction coefficient, *Fz* is the Z-axis value of the force sensor, and *MMF* is the maximum muscle force. As shown in Fig.1B and described above, the sensor sensitivity on the little fingertip during the adaptation session was set to be three times greater than in the baseline session. The default value of *CC* was set to 200, but this value was individually adjusted for the subjects who produced small sensor inputs. The Cursor movement calculated using Eq. 1 was reflected in the 45° directions to the left and right from the vertical axis of the monitor, and the vector created by these defined as the cursor direction (Fig.2A).

**Fig. 2.**
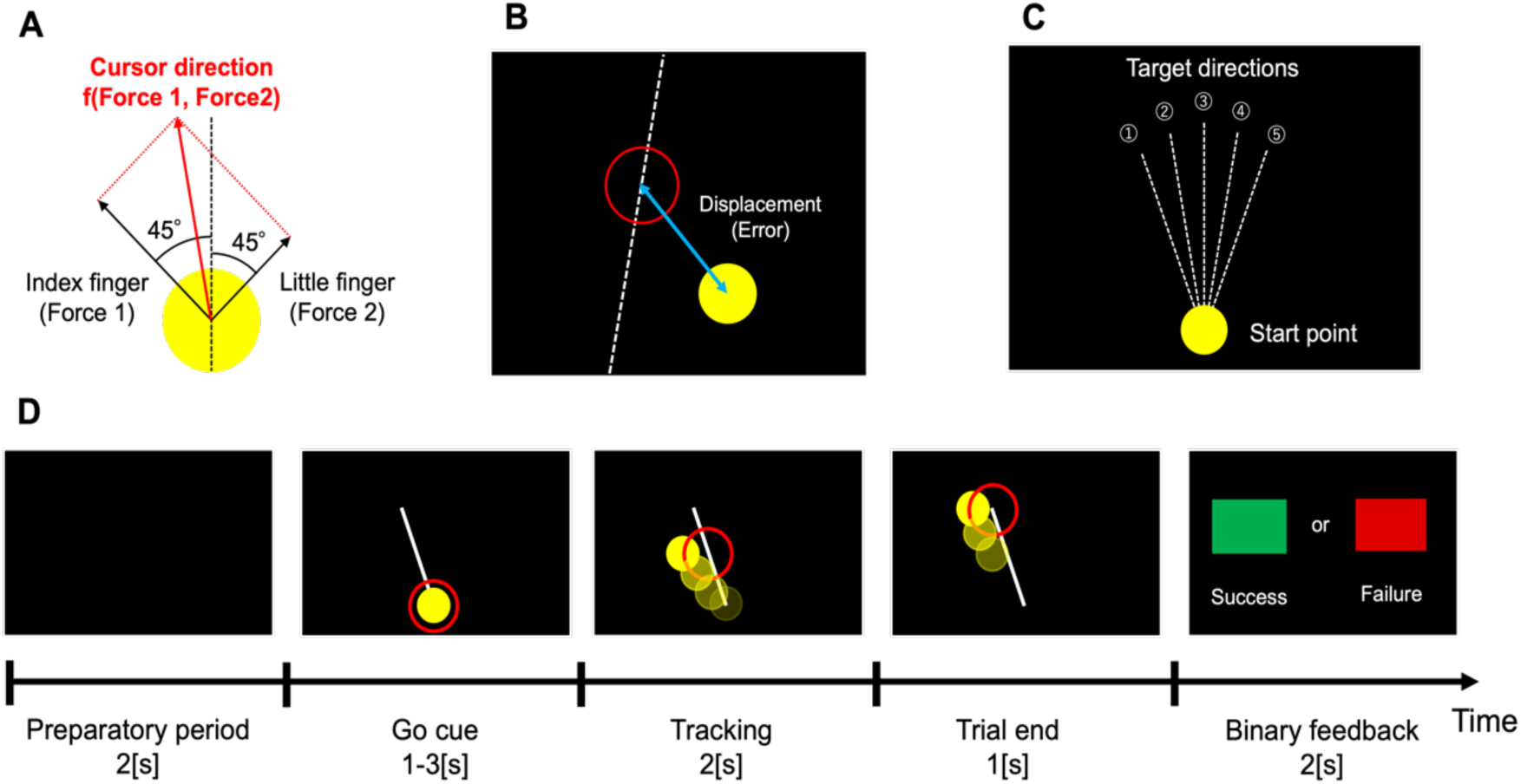
Task structure. **(A)** The cursor direction was determined by a vector based on force inputs from the index and little fingertips of the right hand. **(B)** Task performance was assessed by using the two-dimensional distance between the center of the target and the cursor. **(C)** Target directions were set to five different directions in a randomized order. **(D)** This is the flow of one trial. Due to the time jitter, the timing of the “Go cue” was different for each trial.

On each trial, subjects were instructed to track a moving target on the screen by controlling the cursor as accurately as possible. The target moved straight ahead at a constant speed acceleration about 10.8cm/s in one of five directions (10, 11, 12, 1, and 2-o’clock, Fig.2C) in a randomized order within each session and each direction was set to appear an equal number of times across the individual sessions. Fig.2D illustrates the trial structure. The target circle appeared after a 2-s preparation period, signaling the “Go cue”. The timing of this “Go cue” was jittered, following a normal distribution with μ = 2, σ^2^ = 0.5^2^, a lower limit of 1 s, an upper limit of 3 s. After the “Go cue” appeared, participants tracked the target by controlling the cursor using their force productions. At the end of each trial, they received binary feedback on their performance, green or red squares indicating success or failure, respectively, for 1 s. The luminance of the colors used for the cursor, target, and binary feedback was uniformly adjusted to avoid any biases related to visual perception.

In the present study, we employed different sensor sensitivities in response to the sessions (Fig. 1B). In an initial familiarization session, subjects tracked the moving cursor using a force sensor sensitivity ratio of 1:1 for the index and little fingertips. When performance accuracy exceeded 80%, the diameter of the target was reduced by approximately 0.5 cm (10 pixels) from the default size to minimize the effects of baseline skill on learning proficiency rates. Alternatively, when accuracy was below 80%, the diameter of target was increased by approximately 0.5 cm. After the familiarization session, subjects were asked to complete 50 trials with the normal sensor sensitivity ratio as the baseline session. Following the baseline session, the next 50 trials were assigned to the adaptation session. Throughout this session, subjects had to track the moving cursor using a force sensor sensitivity ratio of 1:3 for the index and little fingertips. This required subjects to learn a new force mapping that combined the use of the index and little fingertips. To prompt exploratory learning processes, we did not provide any verbal instructions about the rate of change in the ratio for the force sensors. EEG data were recorded continuously throughout all task sessions. It should be noted that one left-handed subject also completed the task using the right index and little fingertips.

### Data recording

*Behavioral data.* In accordance with our criterion, only one participant adjusted the correction coefficient in Eq.1 from the default value of 200 to 220. Moreover, three participants reduced the target size from the default size (approximately 6.77 cm) to about 6.20 cm, while seven participants increased it to about 7.33 cm.

*EEG data.* We used a 64-channel EEG amplifier system (actiCHamp, BRAIN PRODUCTS, Germany) to record brain oscillatory activity throughout the experiments. EEG data were continuously recorded from 63 scalp electrodes positioned according to international 10/10 system, using active electrodes embedded in a wearable elastic cap (actiCAP, BRAIN PRODUCTS, Germany), with the ground electrode placed at AFz. The EEG signals were amplified, digitized with 24-bit resolution, and sampled at 1 kHz. During EEG recording, all electrodes were referenced to the right earlobe, while the left earlobe electrode was used for re-referencing during offline data analysis. To monitor horizontal and vertical eye movements, Electrooculography was also recorded from four electrodes placed above and below the left eye and on the left and right sides of both eyes, with a ground electrode placed at the left mastoid. Skin/electrode impedance was kept below 10 kΩ throughout data collection.

## Data analysis

*Behavioral data.* On each trial, performance error was computed using the Euclidean distance between the center of the target (approximately 6.77 cm by default) and the center of the cursor (approximately 5.64 cm) (Fig.2B).

To obtain the performance error, we first calculated the total distance between the center of the cursor and the center of the target during 2-s tracking periods for each trial. This cumulative error was calculated as follow:

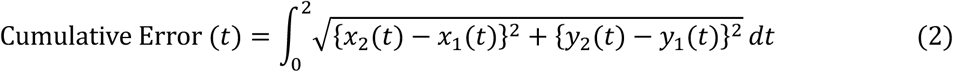

in which (*x_1_(t), y_1_(t)*) represents the cursor’s center coordinates at time *t, (x_2_(t), y_2_(t)*) represents the target’s center coordinates at time *t*. The error for each trial was quantified by integrating the Euclidean distance.

Next, we calculated learning proficiency rates for the new force mapping by exponential data fitting it against the errors from trials of the adaptation session. Exponential fitting was calculated as follow according to a pervious study (Bönstrup et al. 2020):

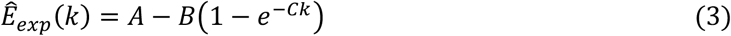

in which *A, B*, and *C* represent the initial value (intercept), the asymptotic value (plateau), and the learning rate (slope), respectively. *k* denotes the trial number. The learning rate *C* was used in the subsequent analysis, where it was treated as parameters indicating individual learning proficiency rates. In exponential fitting, the data for one subject could not be appropriately applied. This dataset was excluded from the EEG data analysis as outlined above.

*EEG data.* EEG analyses were performed using the EEGLAB toolbox (Delorme and Makeig 2004) implemented on MATLAB R2023a (MathWorks, USA) in combination with custom-written code. First, EEG signals were re-referenced to the averaged recordings from electrodes positioned on the left and right earlobe, bandpass filtered (1-50 Hz) and then notched filtered (58-62 Hz) to remove 60 Hz power line noise. Data were epoched to a time window-3200 to 3000 ms that was locked to tracking onset. To remove the artifacts associated with eye movements, eye blinks, and muscle contractions, we applied independent component analysis (ICA). On average, 6.38 components ranging from 2 to 12 components were removed per participant. After performing ICA, epochs containing residual artifacts were detected using an amplitude criterion (±100μ*V*). As mentioned in the Participants section, two subjects had more than 60% of their trials exceeding from the exclusion criterion. Two participants were therefore excluded from subsequent analysis. Finally, to reduce the effects of volume conduction, we applied current source density transformation using the current source density toolbox (version 1.1), which is based on the spherical spline algorithm (Kayser and Tenke 2015). Consequently, 13.6% of the trial data were excluded from the original dataset.

Individual time-frequency power was calculated for each trial within each session using by a Morlet’s wavelet function *w(t, f*):

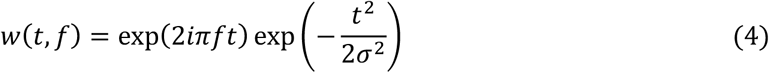

in which *t* denotes time points, *f* represents the center frequency, and σ is the standard deviation of the Gaussian window. The frequency ranged from 1 to 50 Hz in 1-Hz steps, with the number of cycles set to increase from 4 to 10 in linear steps with increasing frequency. This modified wavelet transform was selected to optimize the trade-off between temporal resolution at lower frequencies and stability at higher frequencies (Busch and VanRullen 2010). An event-related desynchronization and synchronization for each trial were normalized using the averaged power between-3200 and-3000 ms from the tracking onset. In this experiment, a timing jitter ranging from 1 to 3 s was set when the visual items were displayed. Due to this, some trials were presented approximately 3 s before the tracking onset. Thus, we selected the time window for baseline correction prior to the item display. We calculated the baseline values for each trial and the baseline values μ_B_(*f, k*) were calculated at each frequency (*f*) and trial (*k*) as follows:

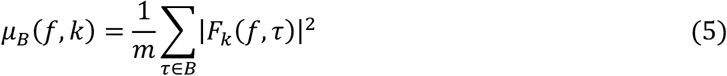

in which τ denotes the time point within the period (*B*:-3200 to-3000 ms) used for baseline correction, *m* is the number of samples during the baseline period, and *F_K_(f, τ*) represents the spectral estimate at frequency f and time τ in trial *k*. The extracted data were converted to decibels after baseline correction as follows:

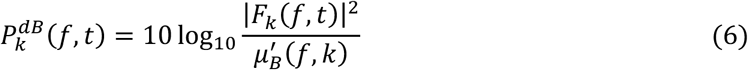

in which *F_K_(f, t)* represents the spectral estimate in the adaptation session. We applied this procedure to both baseline and adaptation trials (Grandchamp and Delorme 2011). The power values of the baseline trials were averaged across trials and the power value during the adaptation was normalized for each trial using the trial-averaged baseline. Subsequently, the number of trials used for averaging was individually determined based on the results of fitting cumulative error using Eq. 3. We believe that this data-driven approach accounted for inter-individual differences in the number of trials needed for error convergence. Specifically, EEG data from the first trial up to the trial where learning was considered complete i.e., the trial on which the fitting curve reached its asymptote, were used for averaging.

Based on our hypothesis and accumulating evidence (Cohen 2011; Cavanagh et al. 2012b; Cohen and Donner 2013; Fine et al. 2017; Jonker et al. 2021), we more focused on FMT as a regional and frequency interest, selecting the FCz electrode, located underneath the ACC. The ACC has been suggested to reflect the necessity for cognitive control (Mitchell et al. 2008; Womelsdorf et al. 2010; Cavanagh and Frank 2014). For the EEG data analysis, the maximum modulation value of FMT power within every 500 ms window was calculated for each trial during motor preparation.

To assess the phase-locking oscillation related to FMT across the trials, inter-trial phase coherence (ITPC) was calculated for each trial, and then was averaged across the trials as follows:

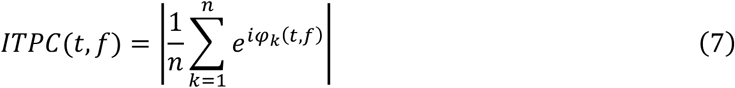

in which *φ_K_(t, f*) represents the phase angle at time *t* and frequency *f* for trial *k*, and *n* denotes the number of trials. A high ITPC is considered that EEG activity at a specified time and frequency is phase-locked to the onset of the “Go cue”.

A previous study reported that the EEG theta phase-locking during motor preparation was consistent exclusively in successful trials, suggesting that the theta phase may contain neural information related to perceptual performance (Tomassini et al. 2017). Building on the previous finding, we investigated how theta phase consistency during motor preparation impacts motor learning proficiency. Specifically, the time point during motor preparation at which ITPC exhibited maximal value was identified and defined as *t_max_* at the individual level.

Subsequently, the mean vector (MV) at *t_max_* was extracted for each frequency bin and denoted as *MV(t_max_, f*).

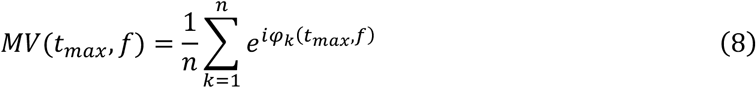

Note that the extracted MV differs in time point across frequencies and participants.

*Statistical analysis.* To address inter-individual differences in motor learning rate and its neural correlates, we investigated the relationship between learning proficiency rates and the maximum modulation value of FMT power during motor preparation. To do so, spearman’s correlation was conducted. To account for outliners of both variable, the skipped correlation method was applied, which controls outliers before computing the correlation (Pernet et al. 2013). An interquartile range (IQR) method was employed for outlier detection. This approach is effective for handling bivariate outliers since correlation analysis is highly sensitive to outliers. Statistical significance was assessed using bootstrapped confidence intervals (CIs) with 1,000 permutations ensuring proper false-positive control. A series of correlation analyses were performed every 500ms during motor preparation starting from 3 s prior to the “Go cue”. To address the multiple comparison problem, Bonferroni-adjusted percentile CIs were calculated (95% CI adjusted for 6 comparisons = 99.166% CI). Correlations were deemed statistically significant if the 99.166% CI did not include zero, ensuring a rigorous assessment of the results.

To examine statistical significance of the relationship between ITPC values during motor preparation and the behavioral measure of learning proficiency rates, a circular-linear correlation coefficient was calculated. This calculation was performed for each frequency within the theta band using MATLAB’s *circ_corrcl* function (Berens 2009). This test assessed the correlation between the cyclic ITPC data and the linear behavioral data, specifically determining whether the data points were distributed along a cylindrical surface.

## Results

*Behavioral results.* Fig.3A shows examples of cursor trajectories from a representative participant when the participant conducted in the first and later trials during the learning condition. Through visual inspection, the deviation in the trajectory decreased as the trials progressed, implying a reduction in performance error and suggesting that motor learning was successfully elicited by the given task. Fig.3B shows the group-averaged performance error normalized to the baseline value throughout the data collection. As expected, the performance error during the first cycle immediately following the onset of the learning condition (i.e., with force perturbation applied) substantially increased compared to the baseline. Fig. 4A shows an example of the performance error along with exponential fittings from a representative participant. As mentioned in the Methods section, learning proficiency rates were represented by the parameter C obtained from the exponential fitting. As shown in Fig. 4B, we observed inter-individual differences in learning proficiency rates and its larger variances.

**Fig. 3.**
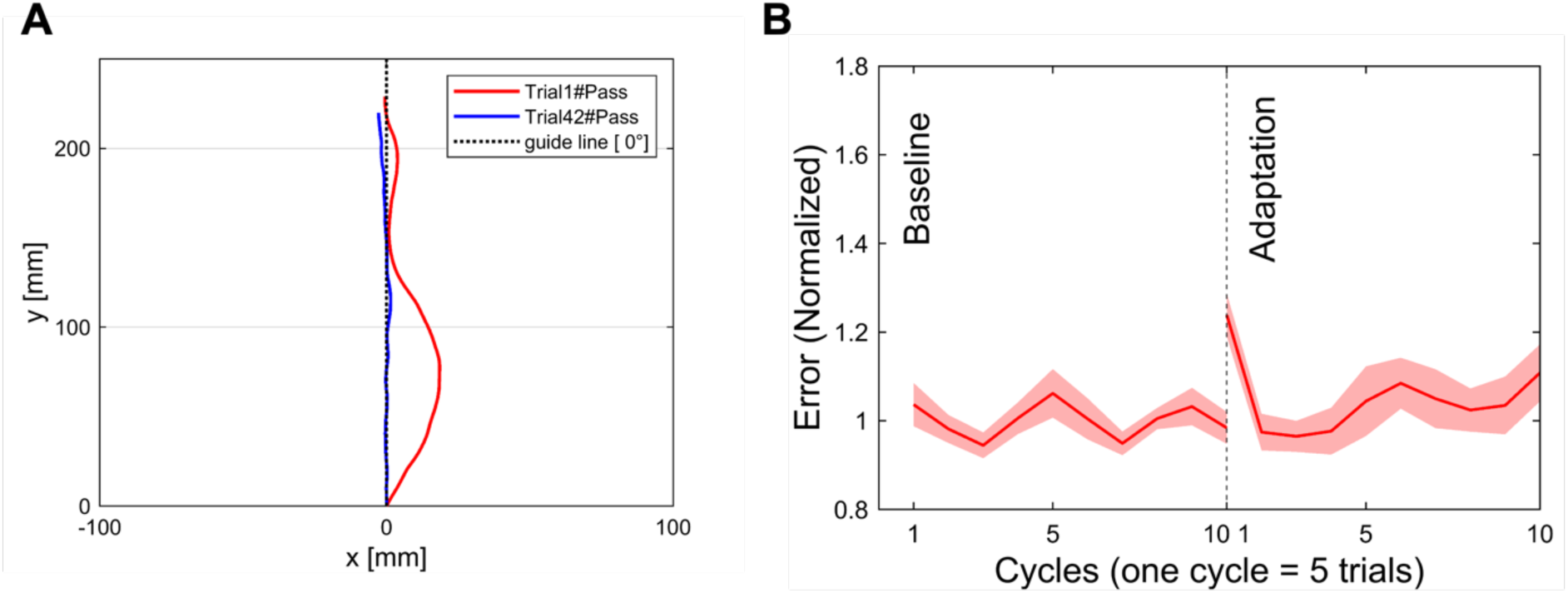
Behavioral Results. **(A)** Examples of cursor trajectories from a representative participant in the early and late learning trials at the 0°target condition. These trajectories exhibit that the tracking accuracy towards the target direction improves in the later trials compared to the early ones. **(B)** Performance errors were averaged across individuals (N=21), with one cycle of five trials. The red shade represents the standard error.

**Fig. 4.**
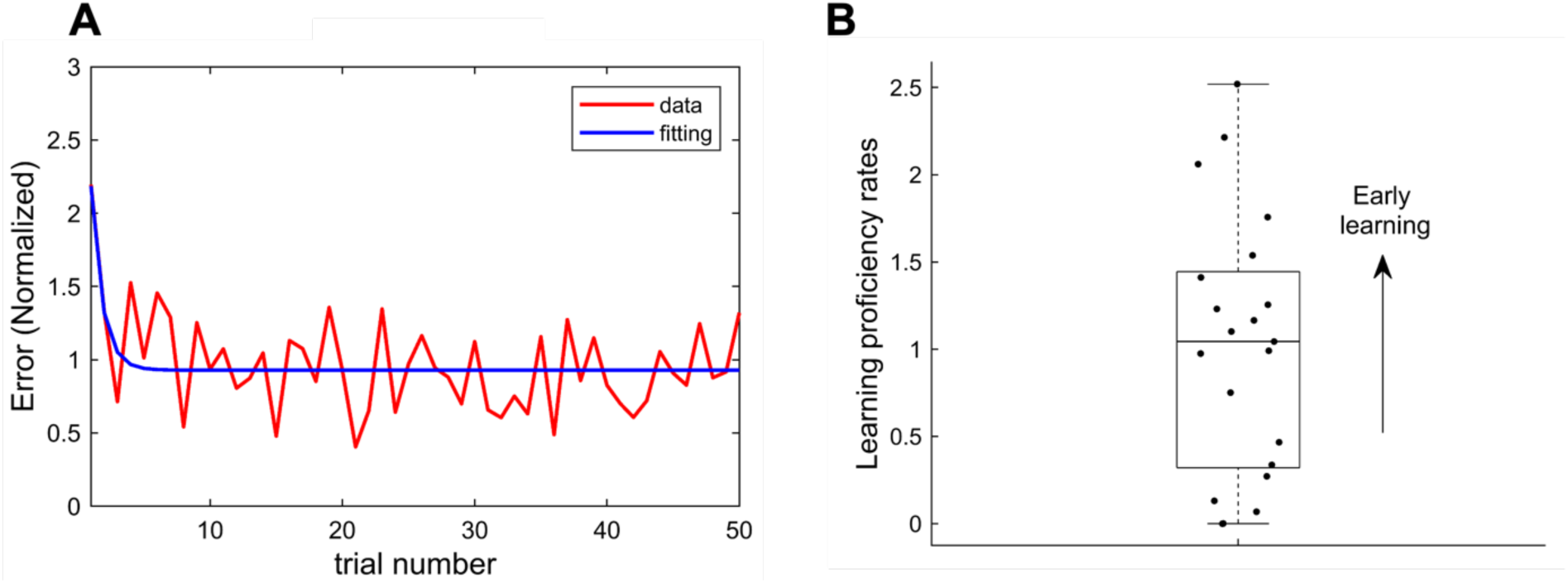
Model fitting of the behavioral data. **(A)** Typical example of model fitting results for the performance errors across the trials of the adaptation session. **(B)** Individual differences in learning proficiency rates. The scatter plot shows the *C* parameter values obtained from the exponential fitting. Each dot represents an individual subject.

*EEG results.* First, the modulations of FMT power were visually compared using data from two representative participants: an early learner (subA, Fig. 5A) and a late learner (subB, Fig. 5B). An individual’s classification as an early or late learner was determined by using their learning proficiency rates, selecting subjects with the highest and lowest learning proficiency rates within the cohort in the present study. For the early learner (Fig. 5A), FMT power exhibited a clear tendency to increase as the tracking onset point (0 s) approached. In contrast, for the late learner (Fig. 5B), minimal changes in FMT power were observed during the same period. Additionally, early learners demonstrated overall greater amplitudes of FMT power modulation compared to late learners. From the perspective of EEG phase in the FMT, the early learner (Fig. 5C) exhibited phase inconsistency (i.e., greater distribution), whereas the late learner tended to be phase-locked across the trials during the adaptation period (Fig. 5D). These indicate that oscillations should not be locked to facilitate motor learning. These observations highlight the importance of further investigating the relationship between FMT and motor learning efficiency in terms of both EEG power and phase.

**Fig. 5.**
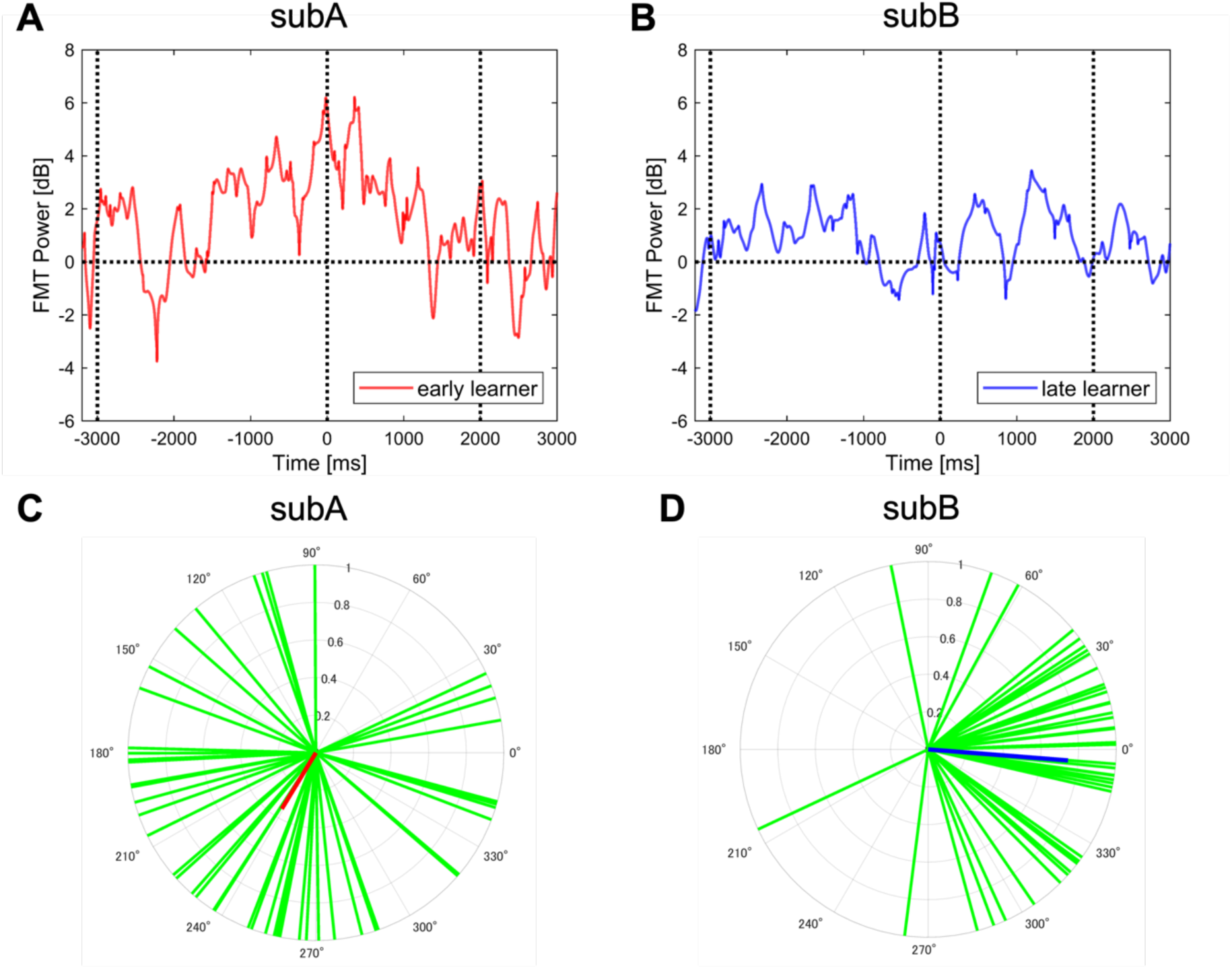
Comparisons of neural characteristics between an early and a late learner. The upper panels illustrate the transition of FMT power averaged across trials during the adaptation session. The vertical dashed lines indicate-3000 ms as the “Go cue” start time, 0 s as tracking onset, and 2000 ms as the end of tracking. The lower panels show the theta phase distribution for each trial, depicted in the green lines. Red (subA) and blue vectors (subB) represent the mean resultant vectors, respectively. Panels (A and C) depict the early motor learner who exhibited maximum learning rate, while panels (B and D) depict the late leaner who exhibited the minimum learning rate within the cohort in the present study.

*Neural correlates.* Spearman’s correlation showed that a significant positive correlation was confirmed between learning proficiency rates and the modulation of FMT power during motor preparation (r = 0.5880, 99.166% CI [0.0114 0.8516], corrected multiple comparisons, Fig. 6). Statistical significance was detected only in the interval immediately preceding the tracking onset, the time-window of-499 to 0 ms. This result indicates that early learners exhibited greater increases in FMT power during the motor preparatory period.

**Fig. 6.**
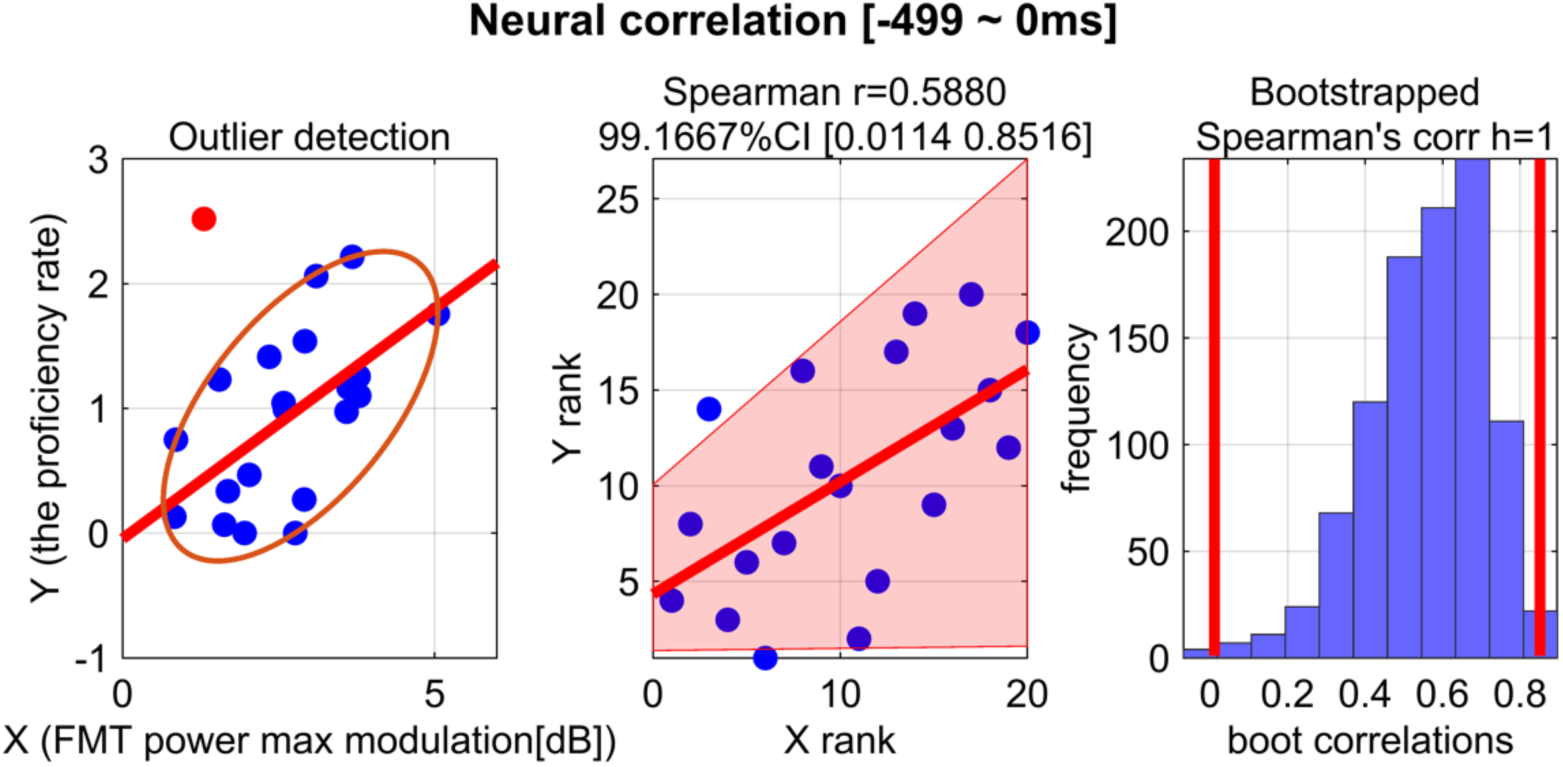
Results of neural correlation between FMT power and learning proficiency rate. Spearman’s correlation was used to test whether a correlation was observed between learning proficiency rates and the modulation of FMT power during motor preparation. Each blue dot in the scatter plots represents an individual. A red dot indicates a participant who exhibited an outliner.

Moreover, to test the statistical significance of the relationship between ITPC value during motor preparation and the learning proficiency rates of behavioral indicators, circular-linear correlation coefficients were calculated. However, no significant correlations were observed across all predefined frequencies (Table 1). This result indicates that no specific EEG phase such as peak, trough, rising slope, or falling slope is associated with being effective for motor learning. In other words, learning effectiveness may emerge from broader neural dynamics.

**Table 1.**
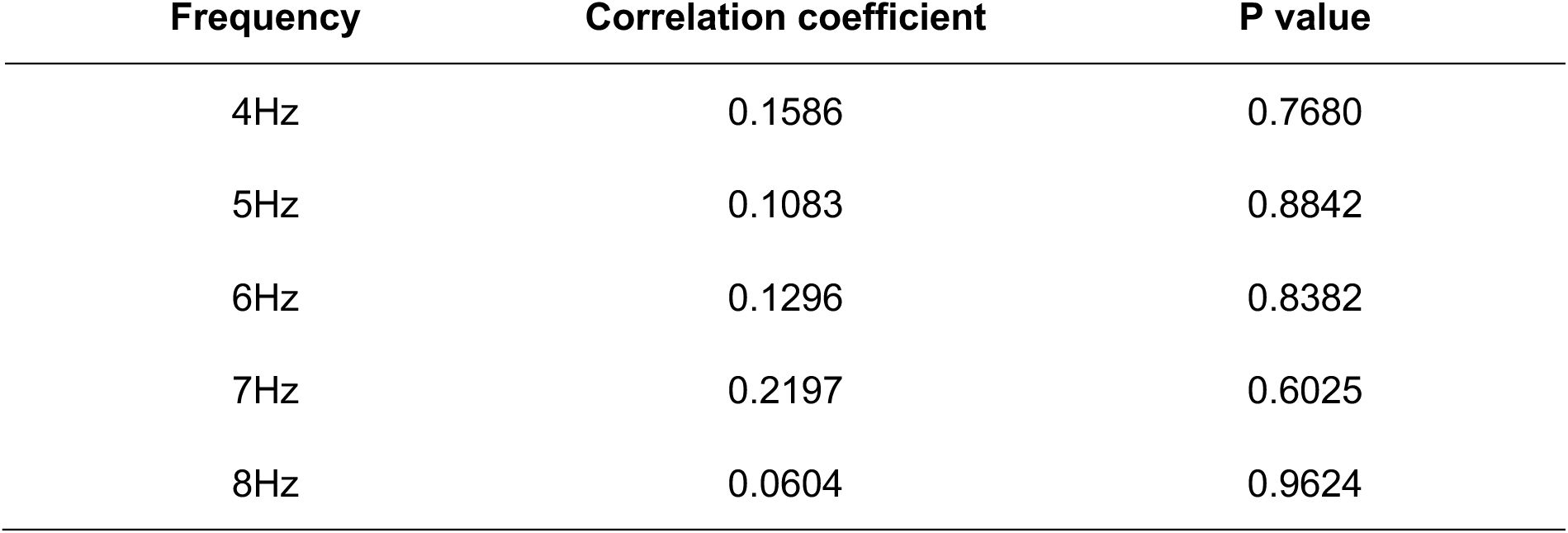
Results of Circular-Linear Correlation Coefficient Analysis.

| Frequency | Correlation coefficient | P value |
| --- | --- | --- |
| 4Hz | 0.1586 | 0.7680 |
| 5Hz | 0.1083 | 0.8842 |
| 6Hz | 0.1296 | 0.8382 |
| 7Hz | 0.2197 | 0.6025 |
| 8Hz | 0.0604 | 0.9624 |

## Discussion

It is a well-established fact that individual differences exist in motor learning. To explore the underlying causes related to brain rhythms, the present study was set up to address whether the modulation of FMT power and theta phase during motor preparation could account for inter-individual differences in learning proficiency rates. We found a significant positive correlation between increased FMT power during motor preparation and learning proficiency rates, whereas no such correlation was observed for EEG theta phase. We provide novel evidence that FMT power, compared to EEG theta phase, serves as one of the neurobiological factors defining motor learning efficiency. Our findings expand the results of the previous studies (Fine et al. 2017; Jonker et al. 2021) and highlight the usefulness of FMT power as an individual factor in defining motor learning efficiency.

Notably, FMT power during the preparatory phase showed a significant positive correlation with the learning proficiency rate. Individuals who promptly completed motor learning exhibited high FMT power. Recently, accumulating evidence suggested preparatory period immediately before action onset plays an important role in motor performance. For example, the selective brain activity related to forthcoming movements begins to occur within the motor related areas (Hasegawa et al. 2017). Cortical network states during the preparatory period impact the feedback control for the skilled motor action (Uehara et al. 2023b). For more direct evidence on learning, neural preparatory activity facilitates motor learning memories (Sun et al. 2022). These findings allow us to posit that changes in preparatory neural activity that accompany motor learning would be linked to changes in motor output and its storing. One of the putative interpretations is that cortical preparatory states may provide the initial condition of the neural dynamics that generate movements afterward and store sensorimotor information to memory. To further clarify this point, we conducted an additional data analysis focused on EEG oscillatory phase and its relationship to motor learning proficiency rates. This is because oscillatory phase has been thought to reflect cyclic fluctuations of a network’s excitability as compared to power changes (Busch et al. 2009). Looking at oscillatory phase attempts to deepen the understanding of neural dynamics during the initial condition, including the phase-resetting model (Kawasaki et al. 2014). Phase-resetting of neural oscillations may serve as timing control of neural information between local and global regions, optimizing responses to upcoming sensory information, reduces the effects of noise and emphasizes signals related to sensory stimuli and actions and predictive encoding and error handling (Makeig et al. 2002; Mazaheri and Jensen 2006; Canavier 2015; Voloh and Womelsdorf 2016). Related to this, phase synchrony (i.e., time-locked to the subsequent action or perception) in the theta frequency band during the preparation for behavior is correlated with perceptual performance (Busch et al. 2009; Busch and VanRullen 2010; Hanslmayr et al. 2013; Tomassini et al. 2017). Furthermore, previous studies have argued whether the phase and amplitude of pre-stimulus oscillatory activity influence the event-related potential in response to sensory stimuli (Barry et al. 2000; Kruglikov and Schiff 2003). Thus, directly comparing EEG power and phase, along with integrative interpretation, is crucial for further understanding the neural basis of individual differences and can provide previously hidden insights. Interestingly, current evidence from human studies suggests that theta oscillations in the medial temporal lobe are associated specifically with binding during memory formation (Clouter et al. 2017). Furthermore, animal studies have demonstrated that synaptic modification related to learning and memory rely on finely tuned timing of the involved neural assemblies in the theta frequency range (Huerta and Lisman 1995; Hasselmo et al. 2002). These pioneering studies have inspired us to evaluate the phase synchronization during movement intentions throughout the learning process. However, as seen in Table 1, there were no significant correlation between preparatory ITPC value (i.e., phase synchronization) across the theta frequency bands and motor learning proficiency rates. Our result suggests that motor learning does not, in fact, rely on the synchronization of neural oscillations in the FMT. Nevertheless, one interpretational caveat is that although non-linear dynamics are widely distributed within the brain, our neural correlation analyses were constrained to linear methods. Our future works will address the remining possibilities including the role of inherent non-linear dynamics in learning function.

In the present study, although we did not perform source localization analysis, the FCz electrode was selected because it is located beneath the ACC, as reported the previous studies (Cavanagh et al. 2012a; Fine et al. 2017; Li et al. 2018; Gawlowska et al. 2018). The ACC is well known for its role in cognitive control (Kerns et al. 2004; Womelsdorf et al. 2010; Myers et al. 2021) and in correcting behavioral responses by reading out of cognitive or motor actions, as well as plans (Lee 2020). In other words, the ACC may serve as a guide for future motor responses. Thus, theta power emerging from the ACC is crucial for determining inter-individual differences in motor learning proficiency rates. This result in line with previous study showing that FMT power is uniquely associated with motor learning, particularly in a context-dependent adaptation model process and the FMT well represents the slow motor learning process in humans (Fine et al. 2017), indicating that FMT can determine the speed of learning acquisition. This brain area also resulted in an increase in activity when changes in learning environment (Behrens et al. 2007). Therefore, rational better understanding is that the ACC and theta oscillations functions such as cognitive control, error correction, and integrating neural information directly impact motor learning process. Nevertheless, addressing physiological underlying the carousal relationship between FMT and motor learning proficiency rates is still open question.

Together, our findings in the present study suggest that although there is a coincidence of local energy distribution within the ACC during the given trial, different neural processes are required in response to the upcoming sensory information with the emerging motor intention. In other words, flexible and high neural activity in the ACC, which is not tied to phase synchronization, may facilitate learning. This compelling interpretation is corroborated by our previous compelling evidence showing that entire cortical network flexibility is one of the factors responsible for skilled motor performance (Uehara et al. 2023b).

The present study has potential limitation. Although this study demonstrated the relationship between the modulation of FMT power and learning proficiency rates, the causal link between them remains unclear. To address this, our future study will extend the present finding to explore causality, such as those employing neurofeedback or non-invasive brain stimulation, e.g., rhythmic transcranial magnetic stimulation (Uehara et al. 2023a). These approaches could modulate FMT power and unmask its causal effect on learning efficiency.

## Conclusion

The present demonstrates the potential of FMT power modulation as a neural indicator that explains individual differences in learning proficiency rates during motor learning. This finding could contribute to the design of personalized learning support systems, including physical education, rehabilitation, and skilled performance in areas such as sports and music. As mentioned above, through causal studies, we will deeply understand the relationship between FMT power and learning efficiency and this process may, in turn, facilitate the development of more effective training methodologies. Understanding individual differences in learning proficiency rates is crucial important in contexts such as education, rehabilitation, and sports training where learning is required.

## Acknowledgments

We thank all laboratory members for their assistance in helping data collection and valuable discussion. We would like to thank the volunteers for participating in this study. This work was supported by JSPS KAKENHI Grant Number JP19K20103 and DAIKO FOUNDATION to K.U.

## Data and code availability

The data and code that produce the findings in this study are available from the corresponding author on reasonable request.

